# Experimental infection of alpacas (*Vicugna pacos*) with Influenza C and D viruses results in subclinical upper respiratory tract disease

**DOI:** 10.1101/2025.07.28.667103

**Authors:** Pauline M van Diemen, Andrew M Ramsay, Helen E Everett, Shellene Hurley, Fabian ZX Lean, Alejandro Nunez, Rebecca Callaway, Adrien Lion, Maria Gaudino, Aurélie Secula, Fatima-Zohra Sikht, Gilles Meyer, Mariette F. Ducatez

**Author notes:** Corresponding authors: Pauline M. van Diemen, Virology Department, Animal and Plant Health Agency (APHA), Weybridge, New Haw, Addlestone, KT15 3NB, UK,; Mariette F. Ducatez, IHAP, 23 Chemin des Capelles, 31076 Toulouse cedex, France. Red Kite Veterinary Consultants, Ltd. 30 Upper High Street, Thame, Oxon, OX9 3EZ.

## Abstract

Influenza D virus (IDV), a new genus within the *Orthomyxoviridae* family, was initially detected in pigs and cattle. IDV is structurally similar to influenza C virus (ICV). Influenza A, C and D viruses all have non-human maintenance hosts and likely circulate in several mammalian species. Camelids, as a reservoir for zoonotic viruses, were not extensively studied until the emergence of the Middle East respiratory syndrome coronavirus (MERS-CoV) in 2012. Antibody responses to both ICV and IDV could be detected in dromedary camels from Kenya but not differentiated, owing to cross-reactivity. It was unclear whether these findings reflected a technical issue or suggested a role for camelids in ICV and IDV ecology. In the present study, therefore, alpacas (*Vicugna pacos*), a camelid species, were experimentally inoculated with ICV (C/Victoria/1/2011) or IDV (D/bovine/France/5920/2014) to assess susceptibility and assess the antibody response. We have demonstrated that alpacas can be experimentally infected with both ICV and IDV with subclinical infection of the upper respiratory tract (URT), suggesting that virus transmission could potentially occur. These findings accord with previous serology results obtained for camelids and indicate a putative role for these species in ICV and IDV ecology.

## Introduction

The newest genus within the Orthomyxoviridae family, influenza D virus (IDV), was first identified in 2011 following the isolation of the prototypal strain, D/swine/Oklahoma/1334/2011, from pigs with influenza-like clinical signs ^1, 2^. IDV is most closely related to influenza C virus (ICV) which causes respiratory infection in humans. ICV and IDV are genetically and structurally distinct from the other viral genera within this family that have epidemic and pandemic potential; influenza A virus (IAV) and influenza B virus (IBV). IAV and IBV have an eight-segmented genome and virions incorporate two envelope glycoproteins, hemagglutinin (HA) and neuraminidase (NA); in contrast ICV and IDV have a seven-segmented genome and a hemagglutinin-esterase-fusion (HEF) envelope glycoprotein (reviewed in ^3, 4^). Despite the structural similarity of ICV and IDV, these two genera only have 50% nucleotide homology and are antigenically distinct ^2^. Additionally, ICV and IDV have different host tropism; ICV is restricted to human populations (reviewed in ^5^), although pigs are susceptible to experimental ICV infection ^6^ whereas IDV has a broader mammalian host range ^7^.

After the first identification in pigs, IDV was isolated from cattle in the United States ^1^ and other countries globally, including France ^8^, Italy ^9^, Northern Ireland ^10^ and Japan ^11^. Genetic data has confirmed the presence of IDV in other countries, including China ^12^, Republic of Ireland ^13^, Luxembourg ^14^, Turkey ^15^, Canada ^16^, South Korea ^17^ and Sweden ^18^. Serological evidence suggests that IDV circulates in other mammalian species including feral swine and wild boar ^19, 20^, Asian buffalo ^21^ and horses ^22^ as well as small ruminants (sheep and goats) and camelids ^23–26^. Additional host species could potentially be exposed to spillover events, as suggested by detection of IDV RNA in aerosols in South East Asian poultry farms ^27^ and human exposure inferred from isolated detections in air samples from communal spaces such as hospitals and airports ^28, 29^. While IDV clinical infection in the human population has not yet been described, serological evidence indicates the potential for occupational exposure ^30,31^. Experimental infection and transmission studies have additionally indicated that mice, guinea pigs and ferrets, a well-recognised model for human influenza, are IDV-permissive animal models ^2, 32, 33^.

In swine, clinical signs linked to IDV infection are observed in the field but experimental infection reportedly results in limited pathology, with viral replication restricted to the upper respiratory tract (URT), although transmission to contact pigs was reported ^2, 6, 34^. Experimental pathogenesis studies of IDV in calves similarly suggested that IDV is a respiratory virus with moderate pathogenicity that can readily be transmitted by direct contact as well as by the airborne route over short distances ^35, 36^. Furthermore, experimental studies indicate that IDV may potentiate lower respiratory tract infections caused by other bovine respiratory disease pathogens during co-infection ^37^. Collectively these observations support the hypothesis that IDV is a predisposing factor for respiratory disease in cattle. Although cattle are a key livestock reservoir for IDV globally, the earliest serological evidence for bovine infection dates back to 2004 ^35^. However, phylogenetic analysis indicates divergence of ICV and IDV between 300 and 1200 years ago ^38^, raising the possibility that another reservoir host exists.

Camelids were not systematically studied as potential reservoirs for zoonotic viruses until the emergence of the Middle East respiratory syndrome coronavirus (MERS-CoV) in 2012. More recently, alpacas were shown to be susceptible to influenza A virus (highly pathogenic influenza A virus of the H5N1 subtype), with infection reported in animals exposed to H5N1 infected poultry on a farm in the USA in May 2024 ^39^. We previously identified anti-ICV and/or anti-IDV antibodies in Kenyan dromedary camels ^24^. Likewise, a second study detected high seroprevalence by hemagglutination inhibition assays in camelid sera from Ethiopia ^25^ and camel sera from Australia, Mongolia, Saudi Arabia, and Nigeria were also highly ICV and IDV seropositive using the same assay in our hands (pers. comm. M. Ducatez in collaboration with Drs. Peiris and Pereradra (University of Hong Kong, Hong Kong SAR)). We were, however, unable to develop an assay to distinguish true positive results from potential cross-reactivity between ICV and IDV antibodies without losing sensitivity. To assess infection, immunological response, and pathogenesis of ICV and IDV in a camelid host species we conducted an experimental infection study in alpacas (*Vicugna pacos*).

## Materials and methods

### Ethics statement

This study was conducted in accordance with UK Home Office regulations under the Animals (Scientific Procedures) Act (ASPA) 1986 under licence PP7633638 and the study number PP7633638-5-001 was reviewed and approved on 30/04/2021 by the Animal and Plant Health Agency’s Animal Welfare and Ethical Review Body (AWERB). Results are reported according to the ARRIVE guidelines ^40^.

### Animals

Seventeen healthy alpacas age 24-37 months were purchased from a UK farm from a herd with bovine tuberculosis free status. Before enrolment in the study, animals were screened for general health status and absence of anti-ICV and anti-IDV antibodies by hemagglutination inhibition (HI) and microneutralization (MN) methods (below), respectively. Throughout the study, all alpacas were housed in animal biocontainment level 2 facilities and received standard pelleted food, with access to hay and water *ad libitum*. To monitor body temperature, all alpacas were implanted with a subcutaneous biothermal IDentiChip (Destron Fearing) in the lower chest caudal axillary region. Alpacas were observed daily for signs of disease and/or welfare impairment and post-challenge, were monitored using a clinical scoring system (**Supplemental Table 1 and 2**) to assess parameters such as alertness, respiration rate, coughing, tachypnea/dyspnea, body temperature, loss of appetite and digestive function.

### Viruses

Influenza C virus strain C/Victoria/1/2011 (ICV) (kindly provided by Richard Webby, St Jude Children’s Research Hospital, Memphis, TN, USA) and influenza D virus strain D/bovine/France/5920/2014 (IDV) (isolated in 2014 by the joined research unit IHAP, Université de Toulouse, INRAE, ENVT, Toulouse, France, and described in ^36^) were propagated in embryonated chicken eggs and on hRT18G cells (ATCC) respectively to produce inoculum stocks. Both strains are well characterized and produced with a minimum number of passages in cell culture to avoid loss of strain fidelity.

### Experimental design

Upon arrival, the alpacas were randomly divided into one group of three control animals (G1) and two groups of seven animals with an average age of 29 months (G2 and G3) with G1 and G2 co-housed. Group 1 (negative control) alpacas were mock-inoculated intranasally (IN) and euthanised after 72h to obtain baseline necropsy samples. The following day, designated 0 days post-inoculation (dpi), alpacas in Groups 2 and 3 were inoculated IN with either ICV (1 x 10^6^ TCID_50_) or IDV (1 x 10^7^ TCID_50_) in a 4 ml volume (2ml/nostril) using MAD300 (Teleflex) delivery of an atomised spray of droplets with 30-100μm diameter.

### Sample collection and storage

Sampling of alpacas was performed following the experimental design in **Figure 1**. Nasal swabs (one per nostril) were obtained before inoculation and daily for the first week starting at 1dpi and every other day during the following week. Daily nasal swabs from each animal were placed together into 2ml of Leibovitz L-15 medium (ThermoFisher Scientific), containing 1% foetal bovine serum (FBS) and 1% penicillin/streptomycin (Gibco, 5000 U/ml) and the swab supernatant was aliquoted and stored at -80°C. Clotted blood samples for serology analysis were obtained using SST vacutainers (BD BioSciences) prior to inoculation and at 3, 7, 14, and 20dpi. Serum was collected and stored at -20°C.

**Figure 1.**
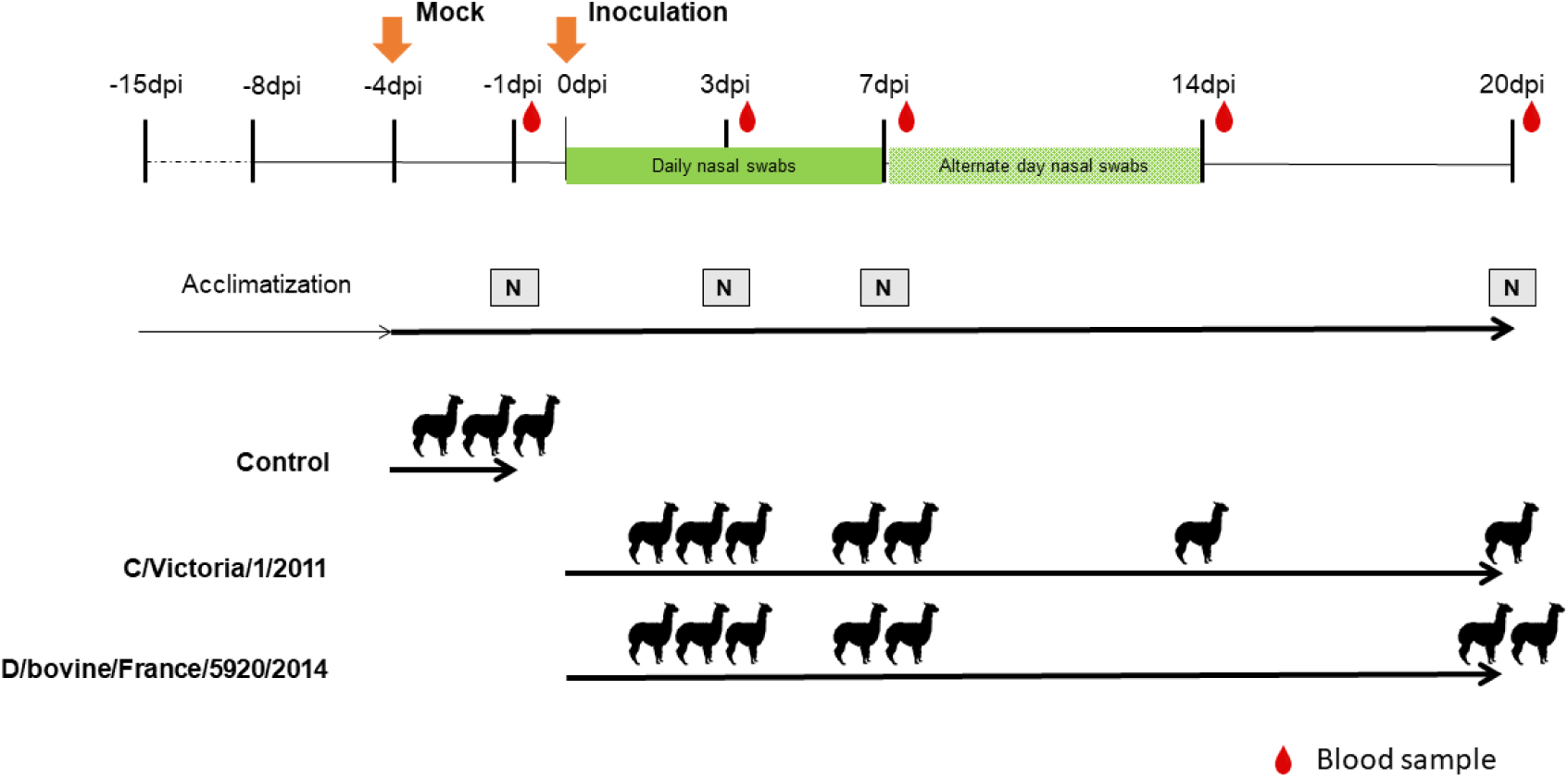
Study design. Alpacas were acclimatized for 10 days. Three control alpacas were mock-inoculated (Mock) intranasally (IN) and humanely euthanised 3 days later for necropsy (N). The following day designated 0 days post-inoculation (dpi), two groups of seven alpacas were inoculated IN with virus strains C/Victoria/1/2011 or D/bovine/France/5920/2014. Clinical signs were monitored daily. Nasal shedding of viral RNA was monitored in nasal swab samples obtained every 1-2 days by real-time RT-qPCR analysis. Blood samples were obtained sequentially on -1-, 3-, 7-, 14- and 20-days post-infection (dpi). Alpacas were humanely euthanised at 3-, 7- and 20-dpi for bronchoalveolar lavage, necropsy (N) and tissue sampling, with one alpaca inoculated with ICV euthanised at 14dpi for welfare reasons.

At defined endpoints on 3, 7 and 20dpi, alpacas were humanely euthanised by overdose of pentabarbitol and a systemic gross examination was carried out by a veterinary pathologist. Necropsy tissues samples were obtained from the nasal conchae, olfactory bulb, trachea, lymph nodes (mediastinal, tracheobronchial, medial retropharyngeal and mesenteric), lung lobes (left and right cranial and caudal, accessory), kidneys, liver, spleen, duodenum, jejunum, ileum, and any other organs showing macroscopic lesions. Additional tissues were sampled for diagnostic pathology in case of underlying pathology not related to experiment infection. Samples were formalin fixed (10% NBF) and stored at room temperature or fresh-frozen in L-15 medium (ThermoFisher Scientific) or preserved in RNAlater (Sigma-Aldrich) and stored at -80°C. Swabs of nasal turbinate and tracheal tissue surfaces taken at necropsy were processed in the same way as nasal swabs. A bronchoalveolar lavage (BAL) sample was obtained from an excised lobe of the left lung using Ca-and Mg-free DPBS (Gibco). The BAL fluid was centrifuged at 1500 rpm for 5 min. at RT and the supernatant aliquoted and stored at -80°C.

### Molecular assays

Viral RNA was extracted from swab suspensions or tissue homogenates using the RNeasy® mini kit (Qiagen). Viral RNA was quantified by real-time RT-qPCR (RRT-qPCR) directed against the PB1-gene of IDV as described ^2^ using the QIAGEN OneStep RT-PCR kit (Qiagen), or against the M42-gene of ICV using iTaq™ Universal SYBR® Green One-Step kit (BioRad) and in-house designed primers (**Supplemental Table 3**). Results are expressed as relative equivalent units (REU) extrapolated from the cycle thresholds by using standard curves generated RNA extracted from the same batch of ICV or IDV, of known TCID_50_ titre, used for inoculation. Although these units measure the amount of viral RNA present and not infectivity, it may be inferred from the linear relationship with the dilution series that they are proportional to the amount of infectious virus present. Viral RNA amounts are expressed according to volume (REU/ml) or weight (REU/g) of sample analysed.

### Histology

Formalin-fixed tissues were processed using standard histological methods. Serial sections of 4 μm thickness were stained with haematoxylin and eosin (H&E) or subjected to *in situ* hybridisation (ISH) using RNAscope®. Twenty pairs of proprietary double Z RNA probes (Advanced Cell Diagnostics, Inc.) were designed against segment 5 (nucleoprotein gene) of Influenza C virus (C/Yamagata/9/2006; NCBI accession LC123782.1; probe catalogue number 1205531-C1; target region 440–1516) and Influenza D virus (D/bovine/France/5920/2014; NCBI accession MG720237.1; probe catalogue number 1205541-C1; target region 190–1131). Both viral genomes were evaluated *in silico* for cross-reactivity against the alpaca reference genome (GCF_000164845.4); no sequence alignment was detected. *In-situ* hybridization (ISH) was performed using the RNAscope® 2.5 HD Brown Detection Kit (Advanced Cell Diagnostics, Inc.) according to the manufacturer’s protocol, as previously described ^41^. Tissue sections were mounted on charged slides, dewaxed in xylene, and rehydrated through graded ethanol. Endogenous peroxidase activity was quenched using RNAscope® hydrogen peroxide for 10 minutes at room temperature.

Antigen retrieval was performed in Target Retrieval Solution for 15 minutes at 100 °C, followed by proteolytic digestion with Protease Plus for 30 minutes at 40 °C. RNA probes were hybridised to tissue sections for 2 hours at 40 °C in a HybEZ™ oven, followed by six rounds of signal amplification using alternating incubations of Hybridise Amp reagents at 40 °C and room temperature (30 and 15 minutes, respectively). Washes with 2× wash buffer were performed for 2 minutes at room temperature between each amplification step. Signal detection was achieved using 3,3′-diaminobenzidine (DAB) chromogen. Sections were counterstained with Mayer’s haematoxylin (Surgipath), dehydrated through graded ethanol and xylene, then mounted with a coverslip using DPX mounting medium (TCS Biosciences Ltd).

### Serology

Sera were treated with receptor destroying enzyme (RDE) (Seika, Japan) and hemadsorbed on packed chicken red blood cells. Antibody titres against ICV strain antigens were measured by an adapted HI assay using 4 hemagglutination units of C/Victoria/1/2011 as an antigen ^24^ A viral microneutralization (MN) assay was used to measure antibody titres against IDV as previously described on swine testis cells (ATCC), using 100 tissue culture infectious doses 50 (TCID_50_) of D/bovine/France/5920/2014 per well and 5 days’ incubation at 37°C and 5% CO2 without TPCK trypsin ^24^. The HI and MN titres are expressed as the reciprocal of the highest dilution of serum where haemagglutination or virus replication was prevented, respectively. Titres strictly above 10 were considered positive ^42^.

## Results

### Clinical and necropsy evaluation

Clinical signs in the alpacas were evaluated daily using a clinical score system (**Supplemental Table 1 and 2**). For most animals, clinical signs remained mild and non-specific, and throughout the study the total score did not exceed 2 out of a maximum possible score of 24. However, elevated body temperature, increased respiratory rate and sporadic diarrhoea were observed in individual alpacas (**Figure 2**) during the infection phase of the study. One alpaca inoculated with C/Victoria/1/2011 (Animal ID: 2019) was euthanised on 14dpi when respiration rate and body temperature were increased over several days, approaching the moderate humane endpoint (**Figure 2**).

**Figure 2.**
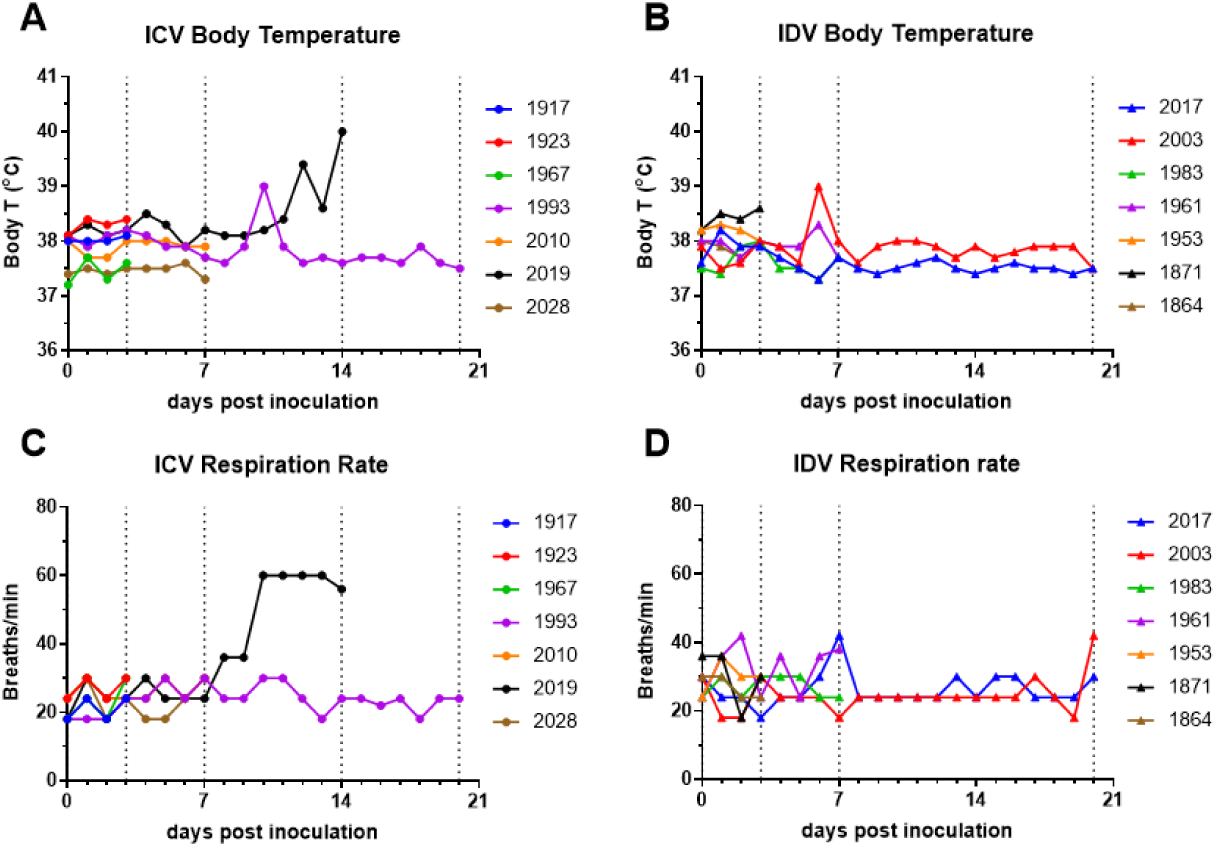
Clinical monitoring. Clinical parameters of body temperature (A, B) monitored by biothermal microchip and respiration rate (C, D) were recorded daily following ICV or IDV inoculation. Vertical dotted lines denote the days on which necropsies were conducted. One ICV-inoculated alpaca (ID 2019) was euthanised on 14dpi when clinical signs approached the moderate humane endpoint.

At necropsy, no apparent gross lesions attributable to viral challenge were observed in the respiratory tract (nasal cavity, trachea, or lungs). However, incidental lesions were noted in three alpacas: one control (Animal ID: 1998) and IDV-inoculated animals euthanised at 3 dpi (Animal ID: 1871), and at 21 dpi (Animal ID: 2003). These included grossly enlarged, firm, white lymph nodes (submandibular, retropharyngeal, tracheobronchial, and mesenteric) with a gritty cut surface, as well as multiple small, firm, white nodules within the hepatic and splenic parenchyma. Similar lesions were not detected in the lungs. Histologically, the lesions were characterised as chronic pyogranulomas, with no detectable bacterial structures on H&E, and no acid-fast bacilli identified on Ziehl–Neelsen staining. The findings were most consistent with a pre-existing chronic infection, unrelated to the experimental viral inoculation.

### ICV and IDV replication

Shedding of ICV and IDV RNA was detected in nasal swabs between 1-7dpi for both viruses and peaked between 2-6dpi (**Figure 3A and B**) suggesting productive infection that was then resolved. Assessment of swabs and tissues obtained at necropsy confirmed the presence of RNA for both viruses in tissues of the upper respiratory tract obtained on 3 and 7dpi (**Figure 3C-F**). The alpaca (Animal ID: 2019), euthanised upon reaching the moderate humane endpoint, had ceased nasal shedding of ICV by 10dpi (**Figure 3A**). Virus was not detected in any tissue of this animal at necropsy.

**Figure 3.**
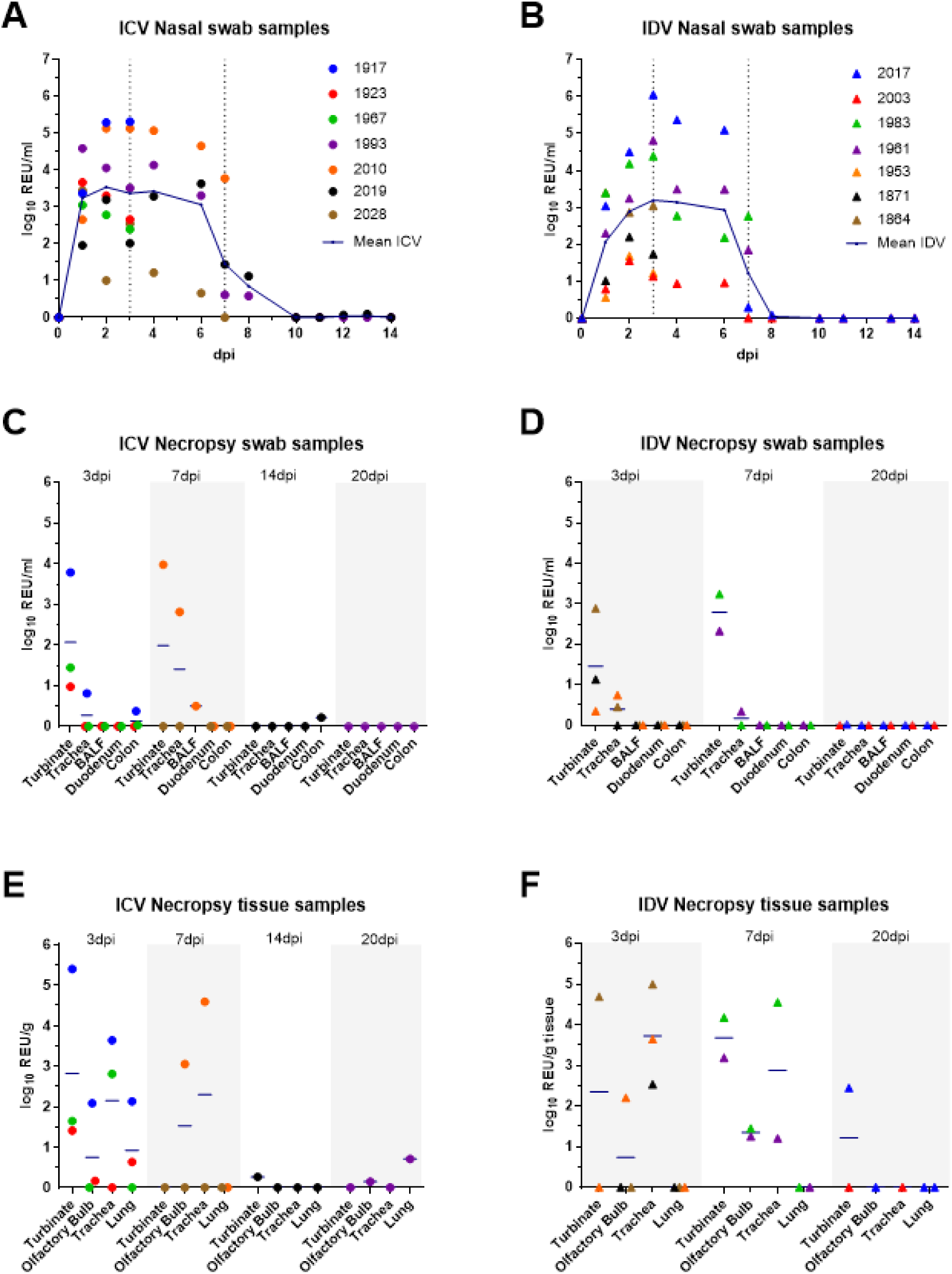
Detection of influenza C virus (ICV) or influenza D virus (IDV) RNA in swab and tissue samples. Viral RNA was monitored in nasal swab samples (A, B) and at necropsy in swab, bronchioalveolar (BAL) fluid (C, D) or tissue samples (E, F) on the indicated days following ICV or IDV inoculation. Viral RNA was evaluated by RRT-qPCR and results are expressed as relative equivalent units (REU) per ml (nasal swabs) or weight (g) tissue. Horizontal lines indicate the mean.

### Microscopic pathology and virus tropism

To evaluate histopathological changes associated with acute infection, tissues from mock-inoculated controls and from animals euthanised at 3 and 7 dpi were examined. No overt epithelial or acute inflammatory lesions were observed in the nasal turbinate or trachea in any animals. In the lungs, there were occasional bronchiolar-associated lymphoid tissue (BALT) in pre-challenge controls. In the ICV-inoculated group, one of three alpacas had mild alveolar leucocytosis in addition to BALT at 3 dpi. At 7 dpi (n=2), one ICV-inoculated alpaca (Animal ID: 2028) had rare intra-bronchial or alveolar neutrophilic exudate and sparse lymphoplasmacytic perivascular cuffs. In the IDV-inoculated group, all animals at 3 dpi (n=3) and 7 dpi (n=2) had infrequent BALT in the lung, similar to that observed in pre-challenge controls. Overall, no consistent or specific microscopic lesions attributable to viral infection were identified, and the presence of BALT in both control and challenged animals likely represents historical immune activation unrelated to the experimental challenge.

RNAscope® in situ hybridisation was performed using probes targeting ICV or IDV segment 5 (nucleoprotein) RNA in respiratory tissues from PCR-positive animals (Influenza C: Animal IDs 1917, 2028, 1967, 1923; Influenza D: 1864, 2028, 1983, 1871, 1953, 1961). Viral RNA (**Figure 4**) was detected exclusively in the olfactory mucosa of the nasal turbinate at 3 dpi (ICV, n=1/3; IDV, n=1/3) and 7 dpi (ICV n=0/2, IDV n=1/2). In one ICV-inoculated alpaca (Animal ID: 1917), approximately 40% of the olfactory mucosal epithelium was positive for viral RNA labelling, with signal localised predominantly to sustentacular cells and occasionally to olfactory neurons. In two IDV-inoculated alpacas (Animal IDs: 1864 and 1961), between 1–10% of the olfactory epithelium was labelled positive, with signal restricted to olfactory neurons. No viral RNA labelling was detected in submucosal glands olfactory nerve fibres or other submucosal connective tissues. Viral RNA localisation did not correspond to any histological lesions.

**Figure 4.**
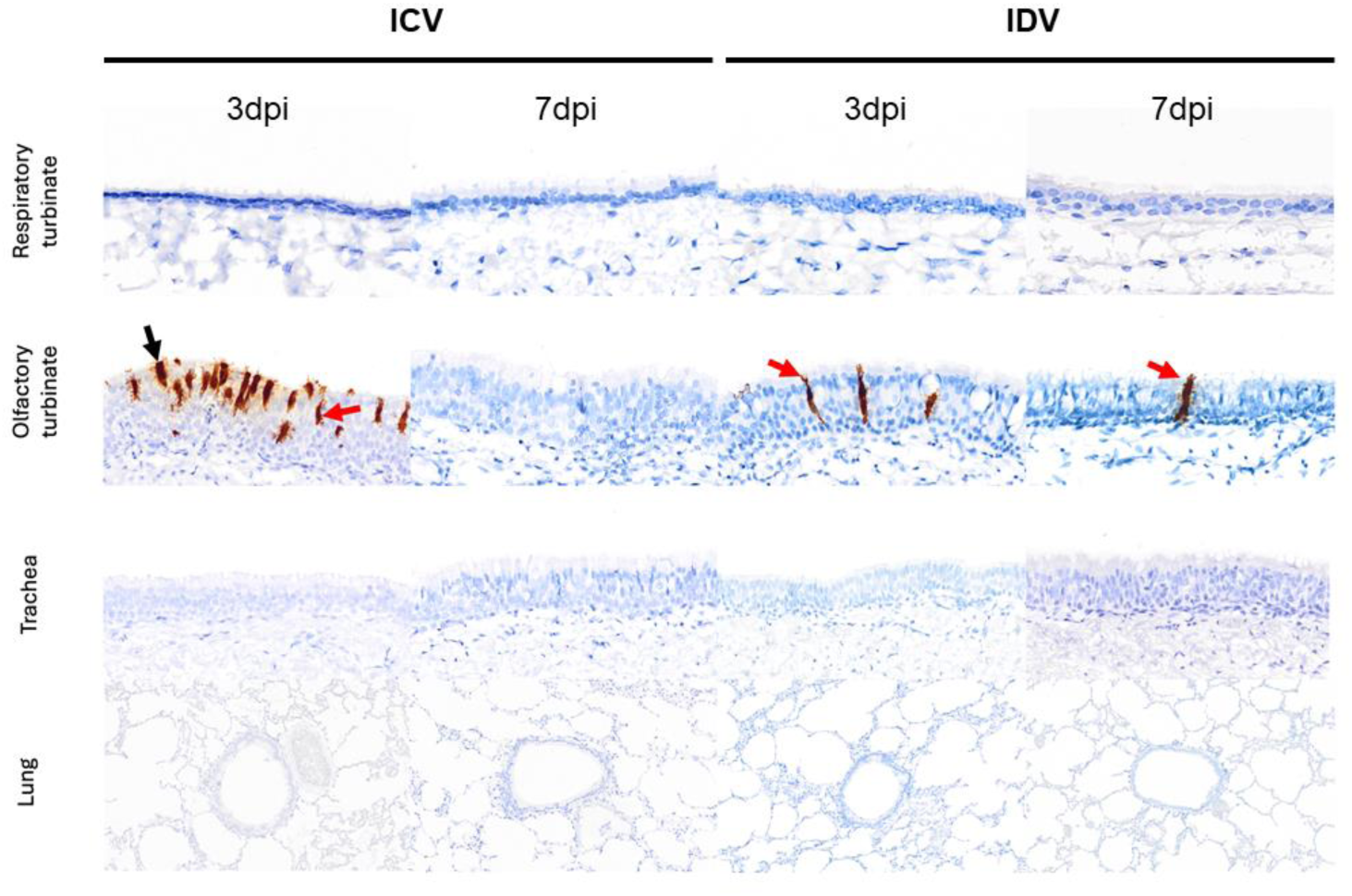
Microscopic detection of influenza C virus (ICV) or influenza D virus (IDV) RNA in the respiratory tract. Viral RNA was visualised by RNAscope® *in situ* hybridisation using a brown chromogen. Necropsy tissues from the respiratory turbinate, olfactory turbinate, trachea and lung obtained at 3- and 7-days post-inoculation (dpi) were examined. Labelling was only detected in the olfactory turbinate, with ICV and IDV RNA detected at 3dpi in sustentacular cells (black arrow) and/or olfactory neurons (red arrow) and at 7dpi IDV RNA could be detected rarely in olfactory neurons. No viral RNA labelling was observed in the respiratory epithelium of the nasal turbinate, trachea, or lung.

### Humoral immune response elicited by ICV and IDV

Specific HI or MN assays were used to assess the humoral responses elicited to ICV or IDV respectively. The immune assays confirmed seroconversion of the ICV and IDV inoculated alpacas from about 14dpi (**Figure 5**). The non-inoculated (control) animals were seronegative for both viruses. Cross-reactions were not observed suggesting ICV inoculated alpacas did not harbour anti-IDV antibodies or *vice versa*.

**Figure 5.**
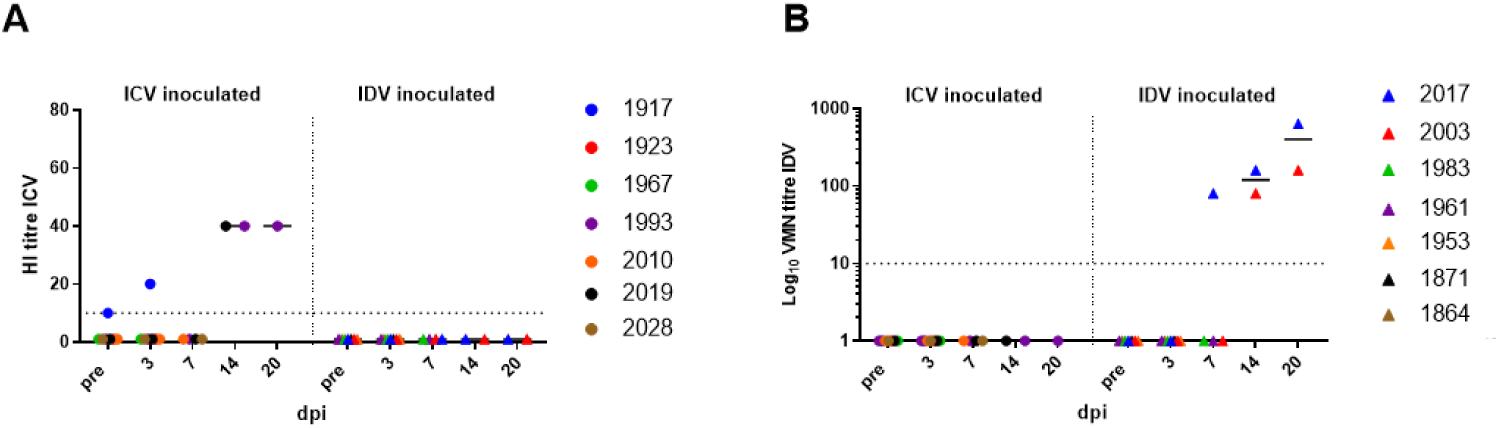
Serological analysis. Serum samples obtained longitudinally pre-inoculation and at 3, 7, 14 and 20 dpi were analysed using (A) Haemagglutination inhibition (HI) and (B) virus microneutralization (VMN) assays. Dotted lines indicate the lower limit of detection for the assay and horizontal line the mean.

The quality of RNA obtained from BAL or nasal swab samples was insufficient to obtain statistically valid transcriptomics results and RNA integrity number ranged between 1.0 and 5.1 and 18S nor 28S peaks could be identified (data not shown). Quantitative RT-PCR results did therefore not allow for a single comparison between controls/infected animals at any day post infection.

## Discussion

The present study was aimed at understanding whether alpaca may be susceptible to ICV and/or IDV infection by considering both clinical signs and viral replication in the respiratory tract. We have demonstrated that alpacas can be experimentally infected with either ICV or IDV, resulting in subclinical infection of the upper airways, with evident virus replication in the nasal turbinate and nasal shedding of viral RNA, suggesting that virus transmission could potentially occur as was already described for MERS-CoV in camelids or SARS-CoV-2 in ferrets ^43^. While olfactory neuronal labelling was occasionally detected, there was no further viral RNA detection in the turbinate submucosa or olfactory bulb and thus no evidence of further dissemination including retrograde neuronal invasion. Inoculated animals seroconverted within two weeks post-inoculation, confirming immune exposure, most likely as a result of ICV and IDV infection. These findings accord with the previous serology results indicating high seroprevalence in camelids and indicate a putative role for camelid species in ICV and IDV ecology. The absence of virus detection in camelids to date may be explained by the very limited clinical signs observed and/or the transient virus shedding: both ICV and IDV RNA could be detected in nasal samples for just a week under the experimental conditions of our study. Additionally, camelids may serve as a subclinical reservoir and maintenance host. They are generally raised outdoors and sometimes with other species (often cloven-hoofed animals). Transmission within camelid populations and between camelids and other species (including cattle) is still unknown and thus warrants further study.

The potential role of ICV and IDV in camelid respiratory disease remains to be elucidated. In cattle, IDV is frequently detected and has been implicated as a contributor to the bovine respiratory disease complex (BRDC), an important multifactorial respiratory disease ^37^. In camelids, respiratory viral infections have been sporadically reported, including those caused by Middle East respiratory syndrome coronavirus (MERS-CoV) and alpaca respiratory syndrome-associated alphacoronavirus, though these typically result in sporadic outbreaks with limited or subclinical disease. ^44–46^ Bacterial agents, including *Actinobacillus vicugnae sp. nov.* ^47^ and *Pasteurella multocida* ^48^, and recently mycotic infection with *Aspergillus fumigatus* ^49^, have been reported in the field. However, it remains unclear whether a camelid respiratory disease complex (RDC) exists, analogous to BRDC in cattle, in which primary viral infections may predispose animals to secondary bacterial or fungal infection.

Perspectives of the present study also rely on a more systematic screen of camelid respiratory tissues (or nasal swabs) to understand etiological agents involved in respiratory disease and their putative interactions as often described in cattle and human population. Cattle are considered to be the main host for IDV and may be the unique reservoir in the absence of any other known host species that may play this role. It is thus interesting to compare IDV pathogenesis in cattle and camelids. The peak of nasal shedding was a bit later (on days 2-8) in cattle than in alpacas (on days 2-6) infected with IDV in experimental conditions and virus shedding was longer (2 and 1 weeks in cattle and alpacas, respectively) ^36^. An inflammatory response was observed in cattle (with mRNA overexpression of pathogen recognition receptors and proinflammatory cytokines and overexpression of the following proteins: TNF-a at d2 and d7, of CXCL10, IL-1-R, and IL-10 at d7, and of IL-2 at d2 and d14) ^36^. In an *in vivo* study on MERS-CoV in alpacas, the authors showed overexpression of genes coding for IFNβ and IFNλ3 (10^2^ fold change range), and to a lesser extent IFNα and IFNλ1 (10 fold change range) could be detected in nasal epithelium on 2 dpi ^43^.

It would be of great interest to study the immune response of camelids post ICV and IDV viruses (MERS-CoV as well as ICV/IDV) trigger similar responses in camelids. RNA conservation is there key. Despite our efforts in storing RNA in the best conditions the RNA quality was insufficient to allow for transcriptomics analyses. Further technological developments may be needed to guarantee absence of degradation of the RNA in camelid BALs and nasal swabs.

The second virus investigated here, ICV, is a pathogen that has been known since the 1940’s. ICV has mainly been studied in human populations, its known reservoir host, where it causes mild respiratory disease mainly in young children and the elderly ^50–53^. ICV has however been detected in dogs, horses, pigs and cattle, again with limited clinical signs associated with the disease ^22, 54–58^. In human as well as in cattle populations, co-infections with ICV and other pathogens have been reported ^57, 59^. Again, the role of ICV in respiratory diseases in its different hosts would need to be further deciphered. While camelids were showed here to be susceptible to ICV as suggested by our previous serology study ^24^, their role in the disease ecology and in virus transmission is still to be clarified.

A limit of the present experimental study is inherent to the small number of animals. Until 7 dpi, few or no clinical signs were recorded, but after that, only 1 or 2 animals remained per group (an ICV inoculated alpaca had to be humanely euthanised), making it difficult to draw meaningful conclusions. In addition, underlying lesions in the lymph nodes could induce innate immune responses and potentially interfere with productive infection (low viral loads were for example detected in one alpaca Animal ID: 1871 (inoculated with IDV) as compared to its counterparts, Figure 3). One can imagine this phenomenon may also be observed in cattle in the field. While Figure 1 (panels A and B) suggests that IDV may replicate to higher titres than ICV, the replication difference may suggest a better adaptation of IDV than ICV to camelids. It may also simply reflect the 2 different inoculation doses (10^7^ and 10^6^ TCID_50_, respectively, due to the difficulty in propagating ICV titres using embryonated chicken eggs (and even more so when attempts to grow stocks on cell lines were carried out)). The differences are in any case difficult to relate to clinical signs while peak of virus titre was in line with peak of respiratory signs in our previous study of IDV pathogenesis in cattle ^24^.

Seroconversion was detected by 14 dpi for both viruses (with an alpaca seropositive for IDV at 7 dpi as well, Figure 5), which is in line with the detection of anti-IDV antibodies in cattle as of 10 dpi ^24^ especially as we did not collect serum between 7 and 14 dpi.

## Conclusions

We have demonstrated that alpacas can be experimentally infected with both ICV and IDV with subclinical infection of the upper respiratory tract, suggesting that virus transmission could potentially occur. These findings accord with the previous serology results obtained for camelids and indicate a putative role for these species in ICV and IDV ecology. Further studies are warranted to assess whether camelids may represent asymptomatic reservoirs for IDV.

## Acknowledgements

The authors would like to acknowledge the staff from Animal Sciences at APHA for excellent animal care. Influenza research at APHA is supported by Defra and the devolved Scottish and Welsh Governments under former and current Research Programmes SE2213 and SE2227 respectively. We thank Richard Webby and Evelyn Stigger (St Jude Children’s Research Hospital, Memphis, TN, USA) for providing strain C/Victoria/1/2011. We thank Ana Moreno (IZSler, Brescia, Italy) for fruitful discussions on the serology assays and Julie Gough (APHA) for assistance with the RNAScope ISH technique. Parts of this study have been presented (presenting author H.E. Everett) at two conferences: as a poster at Options XI for the control of Influenza, 26 - 29 September 2022, Belfast, UK^60^; and as an oral presentation at the 5th ISIRV International Symposium on Neglected Influenza Viruses, 8 - 10 April 2024 Lexington, KY, USA^61^.

## Funding

The research leading to these results has received funding from the European Union’s Horizon 2020 research and innovation programme under grant agreement No 731014 (VetBioNet project) and from the ICRAD-ERA NET co-fund ANR-21-ICRD-0007 “Deciphering the role of influenza D virus in bovine and human respiratory diseases in Europe”.

## Declaration of interest

The authors declare no conflict of interest.

## Data availability

The authors confirm that the data supporting the findings of this study are available within the article and/or its supplementary materials.

## Notes

### Competing Interest Statement

The authors have declared no competing interest.

