## Supplemental tables for "Experimental infection of alpacas (*Vicugna pacos*) with Influenza C and D viruses results in subclinical upper respiratory tract disease"

**Supplemental Table 1. Clinical Scoring guidelines for camelids infected with Influenza viruses.**

| | Parameter | Criteria | Score* |
|---|---|---|---|
| 1 | **Alertness** | Attentive (curious, alert), normal hum (vocalization) | 1 |
| | | Slightly reduced (ears back, avoids handling, self-separation) | 2 |
| | | head down/subdued, reluctant to move, aggressive | 3 |
| | | **Endpoint:** Recumbent, inability to move, inability to access food and water, moaning/teeth grinding | 4 |
| An animal will be euthanized if it has a tendency to stay isolated, shows a delayed response to stimuli (gets up slowly when touched), and/or presents an abnormal posture eg head low for two consecutive days. | | | |
| 2 | **Appetite/leftovers at feeding** | hungry, no increase of left-over feed | 1 |
| | | Slight reduced in take, increase in left over feed | 2 |
| | | Does not eat when fed but tastes food | 3 |
| | | **Endpoint:** Does not eat at all, shows no interest in food, all food is leftover | 4 |
| An animal will be euthanized if it is reluctant to eat at the beginning of the second consecutive day or has weight loss of >12% from heaviest weight recorded. | | | |
| 3 | **Breathing** | Frequency 10 to 30/min, effortless, barely visible chest movement | 1 |
| Prior to approaching | | Frequency 31 to 50/min | 2 |
| | | Frequency 51 to 60/min, flared nostrils | 3 |
| | | **Endpoint:** Frequency >60/min | 4 – see below |
| An animal with respiratory rate >60 the beginning of the second consecutive day will:<br>• be euthanized, If other clinical signs are present<br>• be euthanized at the beginning of the fourth consecutive day, if no other clinical signs are present | | | |
| 4 | **Cough/Dyspnea** | Cough during handling | 1 |
| | | Spontaneous, infrequent | 2 |
| | | Cough and labored breathing | 3 |
| | | **Endpoint:** Open mouth breathing | 4 |
| An animal will be euthanized if it shows no improvement of moderate signs (cough and labored breathing) at the beginning of the second day. | | | |
| 5 | **Temperature (AM)** | 37.5 to 39°C | 1 |
| | | 39.1 to 39.5°C | 2 |
| | | 39.6 to 40°C | 3 |
| | | **Endpoint:** >40°C | 4 – see below |
| An animal with temperature >40°C the beginning of the third consecutive day will:<br>• If other clinical signs are present, be euthanized.<br>• If no other clinical signs are present, be euthanized at the beginning of the fourth consecutive day. | | | |
| 6 | **Digestive system** | Formed regular pellets, normal amount | 1 |
| | | Reduced amount of feces or diarrhea | 2 |
| | | No feces, watery diarrhea, or straining to defecate | 3 |
| | | **Endpoint:** Hemorrhagic diarrhea | 4 |
| An animal will be euthanized if it is passing no feces, watery diarrhea (with no improvement), or straining to defecate at the beginning of the second day. | | | |
| Total clinical score theoretical maximum | | | 24 |

*The individual scores and total score for all 8 parameters are recorded daily per animal. Humane Endpoints are based on these scores

**Supplemental Table 2. Clinical evaluation of daily individual and combined scores and actions to be taken.**

| Clinical assessment | Mild | Moderate | Moderate † | Severe †† |
|---|---|---|---|---|
| Total Clinical* Score | <6 | 7-12 | 13-17 | 18 or greater |
| Maximum for any one parameter* | 1 | 1-2 | 3 for any 2 signs | 3 for any 3 signs<br>4 for any one score |
| **ACTION TO BE TAKEN** | Observe at normal frequency (twice daily) | Inform scientific staff and NVS. Observe at normal frequency with additional check if necessary. | Inform scientific staff and NVS. Observe after 4h. Euthanise if clinical signs worsen or if there is no improvement after 48h. | Inform scientific staff and NVS. Euthanize immediately. |

*Humane Endpoints are based on these scores.
†Moderate severity is recorded if intervention results in improvement and reduction in clinical score after 4h. The humane endpoint is reached, requiring euthanasia, if no improvement occurs after 4h.
††Severe humane endpoint requires euthanasia.

**Supplemental Table 3. RT-qPCR design for detection of influenza C virus (ICV) or influenza D virus (IDV) RNA.**

| | gene | primer | sequence | design | size | GC % | TM | Target size (bp) | RT-PCR kit |
|---|---|---|---|---|---|---|---|---|---|
| ICV | M42 | Fwd | 5'-ACTTGTCAATGGTTTTGTGCTC-3' | this study | 22 | 41 | 56 | 165 | iTaq™ Universal SYBR® Green One-Step (BioRad) |
| | | Rev | 5'-GGAATTGGTGAGTTGTCGGTTT-3' | this study | 22 | 45 | 58 | | |
| IDV | PB1 | Fwd | 5'-GCTGTTTGCAAGTTGATGGG-3' | [1] | 20 | 50 | 57 | 136 | QIAGEN OneStep RT-PCR |
| | | Rev | 5'-TGAAAGCAGGTAACTCCAAGG-3' | [1] | 21 | 48 | 57 | | |
| | | Probe | 5’-FAM-TTCAGGCAAGCACCCGTAGGATT-IBFQ-3’ | [2] | | | | | |
